# Genome wide analysis of YBX1-mediated redistribution of JMJD6 in ER+ breast cancer cells

**DOI:** 10.64898/2026.01.12.698335

**Authors:** Aritra Gupta, Siddharth Bhardwaj, Kartiki V. Desai

**Author notes:** Correspondence to Kartiki V. Desai BRIC-National Institute of Biomedical Genomics P.O. N.S.S Kalyani 741251, India.

## Abstract

The molecular mechanisms by which cancer cells proliferate under diminished ER expression leading to intrinsic and acquired resistance to endocrine therapy remain incompletely understood. Here, we identify a previously unrecognized transcriptional axis centred on JMJD6 and YBX1 that cooperatively reprograms chromatin, redistributes enhancer engagement and sustains tumor proliferation when JMJD6 levels are elevated (JOE cells). Through proximity ligation, ChIP-reChIP assays, KO-based ablation experiments, and genome-wide profiling, we demonstrate that YBX1 is required for JMJD6 enrichment at regulatory sites. ∼20% of JMJD6 and YBX1 peaks physically superimpose, and remaining peaks are contained within a neighbourhood of ±50kb. A shared extended binding site (EBS) containing (i) flanking poly-A/poly-T tracts; (ii) a canonical YBX1 recognition sequence; and (iii) a second complementary motif enriched for zinc-finger proteins defines JMJD6–YBX1 regulated enhancers. Histone modification landscapes in JMJD6 overexpressing cells supported the presence of a transcription permissive chromatin. Further, JMJD6/YBX1 siRNA and overexpression analysis in MCF7 cells identified estrogen and interferon gamma signalling as prominent pathways. The most compelling observation was that ER loss was transcriptional in JOE cells and was accompanied by the emergence of JMJD6–YBX1 enhancer sites at alternate genomic positions relative to canonical ER–JMJD6 sites described in estrogen-stimulated cells. These findings offer a mechanistic insight into earlier reports from our group showing that high JMJD6 with YBX1 suppresses ER expression and this could promote endocrine resistance.

## Introduction

Estrogen/Estrogen Receptor (E2/ER) axis activation is fundamental to gene regulation in normal breast and its prolonged activation is crucial to establish breast cancer. Blocking ER action has greatly reduced mortality in ER+ breast cancers and extended disease-free survival though resistance and non-responsiveness to endocrine therapy remains a clinical challenge (Haque & Desai, 2019). Mechanistically, ER forms a large multiprotein complex with basal transcription factors, mediators, co-activators, histone acetyl transferases, and the epigenetic regulator JMJD6 (Jumanji domain containing protein 6) that allows transcription pause release at ER-target genes (Gao et al., 2018). Earlier, we reported that elevated expression of JMJD6 in ER+ cells, often observed in cancerous but not normal breast tissue, led to a decline in ER levels. JMJD6 orchestrated an alternate gene regulatory network to stimulate ER targets such as MYC, E2F and G2-M transition pathways. This permitted breast cancer proliferation via a JMJD6 dependent but an E2/ER independent axis in these cells (Das et al., 2022). Further, JMJD6 overexpressing cells displayed gene expression profiles similar to Tamoxifen resistant cells. JMJD6 rendered cells insensitive to endocrine therapy and JMJD6 depletion resensitised resistant cells to ER blockers. These observations, in part, explain the observed worse survival outcome in patients bearing tumors with higher JMJD6 expression (Lee et al., 2012).

However, historically since JMJD6 is not a direct DNA binder, to sustain breast cancer proliferation under low ER conditions, it must be recruited at selective ER-target/proliferation genes via interactions with alternate transcription factor(s). Though we initially suspected EZH2 and its interactors to be such a partner, our studies conclusively showed that they acted independent of one another to regulate E2F, cell cycle genes and DREAM targets (Biswas et al., 2020). In the same study, we validated JMJD6 presence by ChIP-PCR, across regulatory regions of multiple genes in these pathways but its recruiting factor remained unknown. In a hunt for such a factor, we utilized mass spectrometry data from JMJD6 antibody mediated pull downs, and using a series of co-immunoprecipitation, co-localisation and domain mapping studies, identified Y-box containing protein 1 (YBX1) as an interacting factor (Gupta et al., 2025). We demonstrated that these proteins positively regulate each other’s expression establishing a positive feed-forward loop and that YBX1 recruits JMJD6 onto the HOTAIR promoter (Gupta et al., 2025). This suggested that JMJD6-YBX1 interaction may have transcriptional consequences (Biswas et al., 2018; Gupta et al., 2025). Prior studies have shown that YBX1 expression affects cell proliferation, motility in breast cancer and alternative splicing in a manner similar to JMJD6 (Dinh et al., 2024). It is known to interact with ER, suppress ER target genes such as pS2 and target ER for degradation (Campbell et al., 2018; Yang et al., 2020). High YBX1 expression is also associated with poorer prognosis and importantly, YBX1 suppresses ER expression to promote endocrine therapy resistance (Habibi et al., 2008; Ito et al., 2012; Stratford et al., 2007). Based on these data, we hypothesized that YBX1 maybe the transcription factor responsible for relocating JMJD6 to alternate chromatin regulatory regions. This study investigates the transcriptional program, genome-wide binding sites and histone landscapes governed by these two proteins and validates this data using ChIP-reChIP, ChIP-PCR and enhancer promoter assays. Mechanistically, our data provides compelling evidence that this interaction redistributes JMJD6 binding on alternate YBX1 directed sites to regulate ER target genes and sustains breast cancer proliferation.

## Materials and Methods

### Cell Culture

MCF7 cells were purchased from American Type Culture Collection (ATCC, VA, USA). The cell lines were grown in Dulbecco’s Modified Eagle’s Medium (DMEM) (GIBCO, USA) with 5% fetal bovine serum (GIBCO, USA) and 1% Penicillin-streptomycin (GIBCO, USA) in humidified 5% CO2 incubator at 37 degrees Celsius. MCF7 cells stably expressing recombinant V5-tagged JMJD6 (JMJD6-V5, referred as JOE cells) and control MCF7 cells carrying only the empty vector (Vec) have been described earlier (Das et al., 2022). Wherever required, transient transfections of JMJD6-V5 and pDEST Myc tagged YBX1 (Addgene, #19878) was carried out as described previously (Gupta et al., 2025).

### Proximity ligation assay (PLA)

5X10^4 cells were plated in coverslips after coating with poly-D-lysine for 24 hours. Fixation was done by using 10% neutral buffer formalin (NBF). Cells were further permeabilized followed by blocking with 10% goat serum for an hour. Overnight probing with primary antibody (JMJD6 and YBX1) was done. After that DUO-LINK kit (Sigma Aldrich) protocol was used. PCR and secondary antibody probing were done from kit provided reagents. Images were captured in high resolution by confocal microscopy. For quantitative analysis we used FijI by Image J and used specific plug in for PLA analysis. We calculated the fold change keeping Vec cells data as 1.

### Transcriptome analysis and validation following perturbation of JMJD6 and YBX1 expression

Data from our previous publication for JOE (GSE211031) and data from YBX1 overexpressing cells (YOE, GSE95299) was analyzed in Geo2R and differentially expressed genes (DEGs) with a 1.5-fold change and adjusted p-value > 0.05 were considered for further analysis. SiRNA mediated knock down (KD) of JMJD6 and YBX1 was performed using reverse transfection method. JMJD6 Si RNA (5’GCUAUGGUGAACACCCUAATT 3’), YBX1 Si RNA (5’ GTTCAATGTAAGGAACGGAT3’) (Ban et al., 2021; Lee et al., 2012) were synthesized (Eurogentec) and for control non-targeting (Scsi) RNA was used (Ambion). RNA-seq analysis was carried out as described in detail in Das et al, 2022. Volcano plots were generated using VolcanoseR, and bubble plots were made using ggPlot2package in Bioconductor R. Validation of RNA seq data was carried out by qRT -PCR analysis using primers listed in Supplementary table 1 and as described in (Gupta et al., 2025). Data for JMJD6si and YBX1si treatment of MDA MB 231 cells were obtained and compared from Biswas et al., 2017 and Lim et al.,2019 respectively.

### Western Blot

Cells were lysed using RIPA buffer, in the presence of 1x Protease inhibitor cocktail (PI) (Sigma Aldrich) and protein extracts were quantified by the BCA method (Pierce, USA). Equal amounts (50 micrograms) of proteins were analysed on SDS-PAGE gels, transferred on to PVDF membrane (Millipore, Germany) and nonspecific sites were blocked by using 5% non-fat milk (Bio-Rad, USA). Primary antibodies were incubated at 4° C overnight and after washing the excess, blots were probed with suitable HRP conjugated secondary antibody for 1 hour at room temperature. Signals were detected using HI-ECL (Bio-Rad, USA) in Chemidoc (Bio-Rad, USA). Antibodies used for Western blots were JMJD6 for total protein (endogenous and exogenous) (PSR, Santacruz, USA, 1:1000), V5 for exogenously expressed JMJD6-V5 (Thermo Scientific, USA, 1:5000), YBX1 total protein (exogenous and endogenous) (Abcam, UK, 1:2000), Myc for exogenously expressed Myc-YBX1 (Sigma Aldrich, USA, 1:2000), β-Actin as internal control (Thermo scientific, USA, 1:5000), Densitometric scanning was done of 3 independent blots at variable exposures.

### Promoter reporter assays

JMJD6/YBX1 binding site regions for VAMP2/AURKB, RAD/BRIX, and IL6, as identified from ChIP, were cloned into the pGL4-TATA vector to assess enhancer activity in luciferase assays. The pGL4-TATA empty vector was used as a control. Previously reported constructs The pRL-TK plasmid was co-transfected as a control in all reactions to determine transfection efficiency across all assays. MCF7, Vec, and JOE cells were transfected with the various constructs using Lipofectamine 3000 and were harvested 48 hours post-transfection. The Dual Luciferase Assay Kit (Promega Corporation, WI, USA) was used to measure luciferase activity. Relative luciferase activity was calculated by comparing the firefly luciferase activity with Renilla luciferase activity. Fold change was calculated relative to the expression in Vec cells, and for the remaining constructs, the RLU of the pGL4-TATA construct was considered as 1.

### Chromatin Immunoprecipitation (ChIP)

Cells were cross linked using 1% formaldehyde solution and reaction was neutralized using 1.375M Glycine. For JMJD6 ChIP, additional cross linker disuccinimidyl gluterate (DSG) was used. For JMJD6 ChIP, V5 antibody and for YBX1, YBX1 antibody was used. For histone marks H3K27Ac (Cell Signaling Technologies, 13027), H3K27me3 (Cell Signaling Technologies, C36B11) and H3K4me1 (Abcam, ab176877) antibodies were used. 10^8^ cells were cross linked, and chromatin was sheared to 300-700 bp fragments in AFA tube using a bath sonicator (Covaris S200). Fragment size was confirmed by D1000 screen tape on Agilent tape station. After pre-clearing for 4 hours antibody pull down was performed overnight with A/G magnetic beads at 4°C followed by protein DNA de-crosslinking. For sequential ChIP, ChIP material from the first pull down was not de-crosslinked, instead, incubation with the partner protein antibody was continued. Isolated DNA was run through PCR purification columns before quantitative PCR with primers for selected chromatin regions (Appendix 1). 10% input DNA was purified prior to ChIP and used for enrichment analysis using the Percentage Input method. For ChIP-sequencing, DNA was further purified and concentrated using Ampure XP beads (Beckman Coulter, Germany). Libraries were constructed using Takara SMART ChIP sequencing kit. Sequencing was performed in NOVA seq (Medgenome, India). 30 million paired end reads (150bpX2) were obtained and after quality control and adapter trimming, alignment with human genome GrCh38 was performed using Bowtie 2. MACS2 was used to call peaks. For, histone marks only broad peaks were considered. Further visualization and analysis were done by ChIPseeker package in Bioconductor (Yu et al., 2012, 2015). Deeptools package was used in Python to make the bigWig files for display in the UCSC genome browser. For ENCODE ChIP seq data of histone marks: H3k27Ac, H3K4me3, H3K4me1 in MCF7 cells were downloaded and similar analysis pipeline was followed.

### JMJD6/YBX1 knock out

JMJD6 and YBX1 knock outs were carried out by using CRISPR. Guide RNA (sgRNA) was designed by using a tool named CRISPOR. Further prioritization of sgRNA was made based on their least off target effect and sequences were cloned in e-sp-Cas9-2a-GFP vector. Cells with knockout of YBX1 (YKO) had shown a lethal phenotype hence we used transient KO of cells for all our experiments.

### Analysis of patient samples

26 breast cancer punch biopsies were used to perform RNA-seq analysis and gene expression as validated by RT-PCR. Institutional review board approval and informed consent was sought for this study (NIBMG/20/2018-19). Protocols used were similar to those described for breast cancer cell lines above. Firehose legacy data was used to download gene-specific TPM counts. Gene expression correlation analysis was performed using KM-plotter online (https://tnmplot.com/analysis/) using RNA-seq breast cancer data.

## Data Availability

Transcriptomic and ChIP-seq data generated in this paper is deposited in GEO omnibus. (GSE211031 and GSE296699 respectively)

## Statistical analysis

All the experiments are done three times; graphs are made by Graph pad prism. Statistical analysis was done by R. Comparisons between two groups were performed using student’s t-test and data was considered statistically significant at p ≤ 0.05. Significance in all figures is indicated as follows: * p ≤ 0.05, **p ≤ 0.01, ***p ≤ 0.001 and ****p ≤ 0.0001.

## Results

### JMJD6 and YBX1 mediated gene regulation

Our recent study showed that YBX1 recruited JMJD6 to the HOTAIR promoter suggesting that their interaction may have transcriptomic consequences (Gupta et al., 2025). Since YBX1 is localized to both nuclear and cytoplasmic cellular compartments, proximity ligation assay was carried out and as shown in Figure 1 (A and B), a robust 12-fold increase in nuclear signal was evident in JOE cells when compared to Vec cells. This indicated nuclear presence of JMJD6 and YBX1 proteins within 40 nm vicinity. To determine their effect on gene transcription, both siRNA treatment and over expression studies were undertaken in MCF7 cells. We have previously documented gene expression profiles of JOE cells (GSE211031) (Das et al., 2022) and we obtained YBX1 overexpression profiles (YOE) from GSE 95299. Using GEO2R analysis, 1886 (780 downregulated; 1106 upregulated) were found in JOE and YOE cells had 3379 (1507 downregulated; 1872 upregulated) differentially expressed genes (DEGs) respectively (*p* value <0.05, FC=1.5). 420 genes overlapped between JOE and YOE cells (Figure 1C). In parallel, RNA-seq analysis of MCF7 cells treated with JMJD6 and YBX1 siRNAs found 1401 and 2421 DEGs respectively (*p* value 0.05, FC=1.5; Figure 1D and 1E). JMJD6si treatment upregulated 774 and suppressed 627 genes whereas YBX1si affected 1163 and 1258 genes respectively (*p* value 0.05, FC=1.5; Figure 1B). 981 genes were regulated by both JMJD6 and YBX1 (Figure 1F). Gene lists are available in Supplementary file 1. Since both JMJD6 and YBX1 showed high expression in MDA MB231 cells and high positive correlation in gene expression across the TCGA ER-negative Breast cancer dataset (Supplementary figure 2A), DEGs from siJMJD6 and siYBX1 treated cells were identified. Data revealed 887 commonly regulated DEGs that involved cancer metabolism, DNA repair and cell cycle (Supplementary figure 2B). Ten DEGs from the MCF7 OE and SI set corresponding to various patterns of regulation across the 4 conditions mentioned above were validated by qRT-PCR analysis in independent sets of transiently transfected MCF7 cells (Supplementary Figure 1). Six genes, *AURKA*, *AURKB*, *IGF2BP3*, *DNAJC21*, *SIRT4* and *BRIX* were chosen since we had reported JMJD6 binding sites (JBS) in their regulatory regions (Biswas et al., 2020). Except SIRT4, all others showed increased expression in cells transiently transfected with expression constructs and siRNAs reversed their expression. In addition, 4 genes with unknown JBS binding site status, *DEGS2*, *SCART1*, *ABHD2* and *IRF1* were used to validate siRNA data. Curiously, overexpression of both JMJD6 and YBX1 suppressed several genes at severely high fold changes but induced genes at a moderate fold change. Conversely, this range of regulation was reversed in the siRNA data. Genes were induced at high manifold but suppressed at lower fold changes. These observations indicate a post-transcriptional function for JMJD6 and YBX1 in RNA stability which requires further exploration.

**Figure 1:**
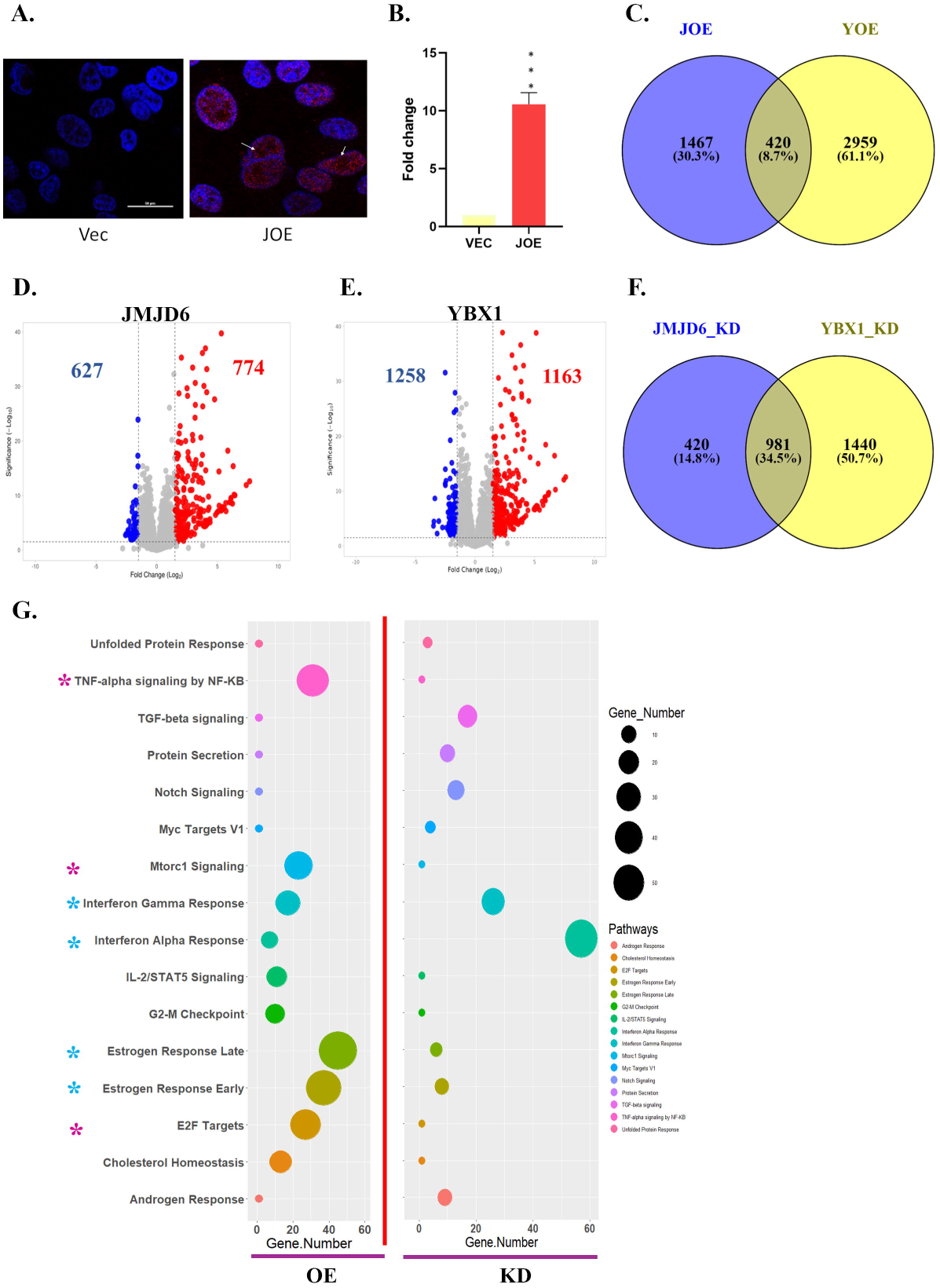
Transcriptional regulation by JMJD6 and YBX1. A) Proximity ligation assay (red dots) showing interaction between the two proteins, nucleus is stained by DAPI (blue); B) Bar graph depicting quantified values of PLA images; C) Venn diagram showing commonly regulated transcriptional targets in JOE and YOE cells; D.-E) Volcano plot of siJMJD6 and siYBX1treated cells respectively. Overexpressed genes are shown in red, and downregulated genes are shown in blue, unaltered genes are in grey colour; F) Venn diagram showing commonly regulated transcriptional targets in siRNA treated MCF7 cells; G) Pathway enrichment analysis of common transcriptional targets after over expression and knockdown. Blue asterisks denote pathways similarly regulated by both proteins, and red asterisks denote contrasting pathway.

Next, pathway enrichment analysis was carried out to determine the most likely cellular functions that are impacted by both JMJD6 and YBX1 (Figure 1G). Interestingly, Estrogen response early, Estrogen response late, Interferon alpha, Interferon gamma regulated genes were regulated in both the OE and SI set of genes whereas the E2F, MTORC1 and TNF-alpha signaling genes were restricted to the OE gene set (Figure 1G, cyan and pink asterix respectively). Clearly, the noteworthy genes regulated by both proteins were either ER-targets and/or immune related genes.

### Binding of JMJD6 to its DNA binding sites (JBS) may be YBX1 dependent

Encouraged by the alteration in gene expression of candidate genes bearing JBS and the HOTAIR data published earlier, we asked if YBX1 interacted with previously validated JBS. 20 JBS sites detected in HeLa cells (HeLa JBS) were tested and 13 of them were validated in both JOE and MDA MB 231 cells by ChIP-PCR analysis (designated as BrCa JBS) (Biswas et al., 2020). To determine if YBX1 co-occupied these sites, YBX1 ChIP was performed and PCR primers specific for BrCa JBS sites were used to estimate YBX1 enrichment. Three non-binding sites (NBS) were included as control. A positive enrichment was observed for all BrCa JBS using YBX1 ChIP material, suggesting that YBX1 may co-occupy JMJD6 bound regions (Figure 2A). Similar experiments using YBX1 ChIP material from MDA MB 231 cells showed YBX1 occupancy at JBS sites (Supplementary figure 2, C and D). To confirm co-occupancy, ChIP-reChIP assays were performed, that is, JMJD6 ChIP material was used as an input to perform a second ChIP with YBX1 antibody and vice versa. As shown in Figure 2 (B and C), JMJD6-re-ChIP assay validated 10/13 sites whereas YBX1-reChIP validated 7/13 sites. This difference in validation could be attributed to technical variation in crosslinking since dual crosslinking agents (DSG and formaldehyde) were used for JMJD6 versus single agent (only formaldehyde) was used for YBX1. The two antibodies used for ChIP may also have differential efficiencies in pulling down DNA and different amounts of DNA may be available for the second (re)ChIP. To circumvent these technical challenges, YBX1 was knocked out in JOE cells via CRISPR knock out strategy (Figure 2D, Inset). In these YKO cells, a significant loss in JMJD6 enrichment was recorded for all 13 BrCa sites. These observations substantiate the importance of YBX1 in recruiting JMJD6 to DNA.

**Figure 2:**
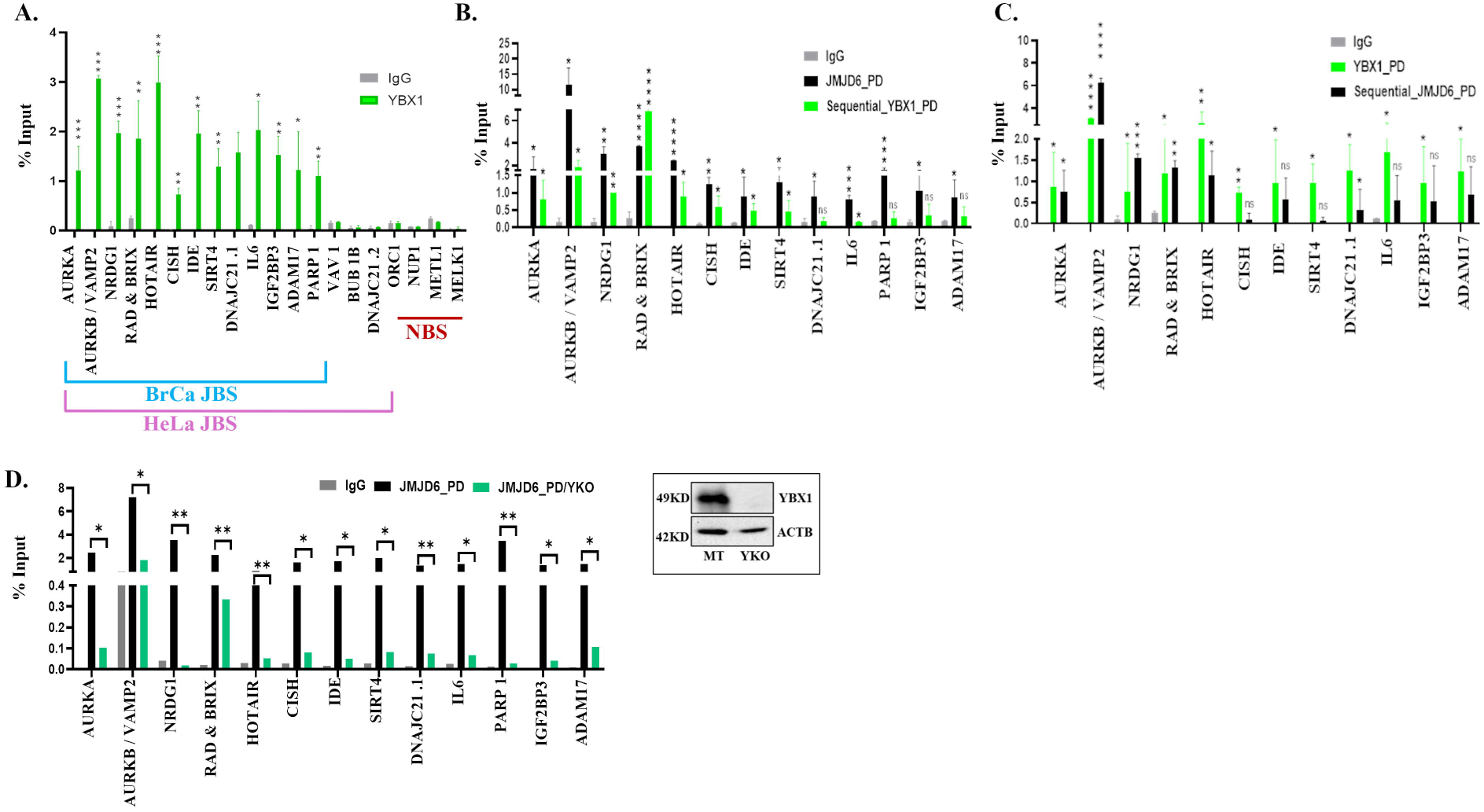
YBX1 binds and recruits JMJD6 to its binding sites. A) YBX1 ChIP and PCR in sites previously identified as occupied by JMJD6 in JOE and HeLa cells, B) ChIP re-ChIP assays and enrichment of at JBS (JMJD6 ChIP followed by YBX1) and C) Vice-versa (YBX1 ChIP followed by JMJD6); D) JMD6 ChIP before and after CRISPR mediated YBX1 knock-out in JOE cells. Immunoblot validates loss of YBX1 expression (inset).

### Genome-wide binding site analysis and motif determination

To assess the extent of co-occupancy in genome wide binding sites of JMJD6 and YBX1, ChIP sequencing was performed in JOE cells. 30 million reads (paired-end, 150bp) were obtained for ChIP-seq libraries, and these were analyzed as described in Methods. After peak calling, we filtered the data based on adjusted *p* value (Benjamini Hochberg corrected, p<0.01) and a fold change cutoff over input, of 4 (JMJD6) and 3 (YBX1) was used to obtain 54719 peaks and 42497 peaks for JMJD6 and YBX1 respectively (GSE296699). Peaks obtained were annotated and distribution of JMJD6 (JBS) and YBX1 peaks (YBS) relative to TSS was conducted using ChIPSeeker package. Both proteins displayed strikingly similar profiles of peak distribution. Very few (about 20%) of total peaks for both proteins were found in the promoter region, and peak frequency increased away from TSS (Figure 3A). More than 50% of the peaks were found 10-100kb away from TSS (Figure 3B). Finer peak annotation revealed that maximum peaks were present in the intronic and distal intergenic regions for the two factors (Figure 3C). However, similar overall distribution is no confirmation of co-occupancy. To test if JMJD6 and YBX1 co-bound similar genomic locations, overlap enrichment analysis was conducted using the ChIPSeeker package (p<0.001, 1000 iterations). Of the total sites, 9733 (∼20%) of the chromosomal regions were superimposed on one another suggesting that they were co-occupied by both proteins. Figure 3D shows the distribution of YBX1 sites very close to the peak summit of 9733 JBS. For the remaining sites, it was possible that JMJD6 and YBX1 interact via DNA looping interactions. To assess this, we determined the peak summit of JBS and extended the genomic co-ordinates on either side in steps of 5-10 kb and scanned for overlapping YBX1 binding regions. Almost all YBX1 sites encountered within ±50kb region of JBS (Figure 3E).

**Figure 3:**
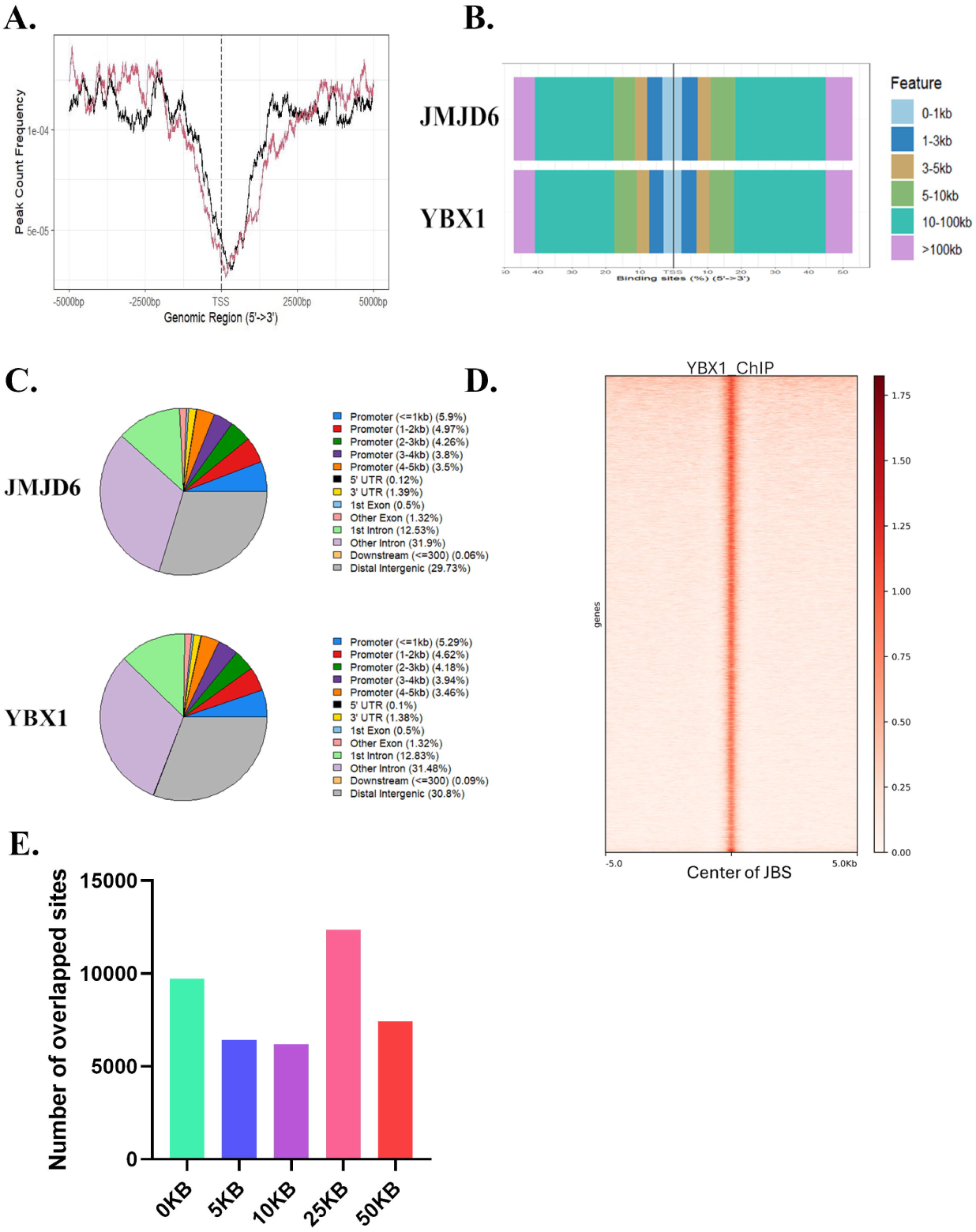
Characterization of genome wide binding sites of JMJD6 and YBX1. A) Peak frequency of JMJD6 and YBX1 around TSS; B) Genomic Features and distance of binding sites from TSS. C) Annotation of binding sites for regulatory features. D) Heatmap of YBX1 peaks relative to the centre of JMJD6 peak summits; E) Overlap enrichment analysis to detect YBX1 binding in the neighbourhood of JMJD6 binding sites.

To date, no specific motif has been defined for JMJD6 binding since it is a secondary DNA binder and uses the DNA binding ability of interacting transcription factors for recruitment into transcriptional complexes. YBX1 on the other hand is a DNA binding factor. It was first shown to bind Y/CCAAT enhancer sequences but analysis of ChIP data from various cell lines has identified 5 additional binding motifs(Dolfini & Mantovani, 2013). Due to this lack of consensus, we used MEME-ChIP suite to scan a maximum of 6 possible motifs in 9733 sites from individual sequence data of JBS and YBS chromosomal locations. The average input sequence was 270 nucleotides. As expected, motifs obtained were very similar for the two proteins since they were predicted from overlapping sites (Figure 4A). The first observation was that motifs were arranged in a manner such that JMJD6 ChIP sites were flanked by a homopolymeric poly-A stretch and YBX1 were flanked by poly-T sequence (Figure 4A and B, Motif 1). The second motif for JBS and YBS displayed complete sequence complementarity and also contained a known YBX1 motif (Figure 4A and B, Motif 2; YBX1 site marked by an oval). The third motif was identical in terms of both sequence and directionality in both JMJD6 and YBX1 derived binding sites but different from Motif 2. The remaining motifs were permutations and combinations of complementary and reversed sequences identified in the first 3 motifs (data not shown). Display of motifs using MEME for select binding sites revealed that multiple motif combinations in the same binding region suggesting a multiprotein complex maybe seeded by JMJD6-YBX1 binding (data not shown). We wondered if this polyA/T stretch was a consistent feature in previously identified Hela JBS sites. We revisited the BrCa JBS which are a subset of Hela JBS and found that all those sites also displayed a combination of Motif 1-3 (data not shown). This made us redefine the specific motifs 2-3 plus Motif 1 as an extended binding site (EBS) interacting with JMJD6 and YBX1. Further, in Motif 2, the additional sequence apart from the YBX1 sites were present and to find possible enriched TFs, TOMTOM analysis was conducted. This analysis identified a series of Zinc finger proteins as possible interactors, suggesting that more undiscovered factors could be components of the EBS. Further analysis using JASPER indicated that ZNF460 (Motif 2) and/or ZNF213 (Motif 3) were the most likely factors that may interact with these sites in a sequence specific manner (Supplementary Figure 3). Interestingly, non-overlapping sites (+/-) 5-50 kb displayed polyA half Motif 2 for both JMJD6 and YBX1 sites and the ZNF460 motif. Motif 3 and the ZNF213 sequence was found for JMJD6 but not encountered in YBX1 sites away from the JBS peak. JMJD6 likely interacted with Motif 3 sequence but published data did not identify any known interaction between JMJD6 and ZNF213. Since half motifs were discovered in non-overlapping sites, it is likely that JMJD6 may loop to YBX1 over these chromosomal locations and Hi-C data could resolve this issue.

**Figure 4:**
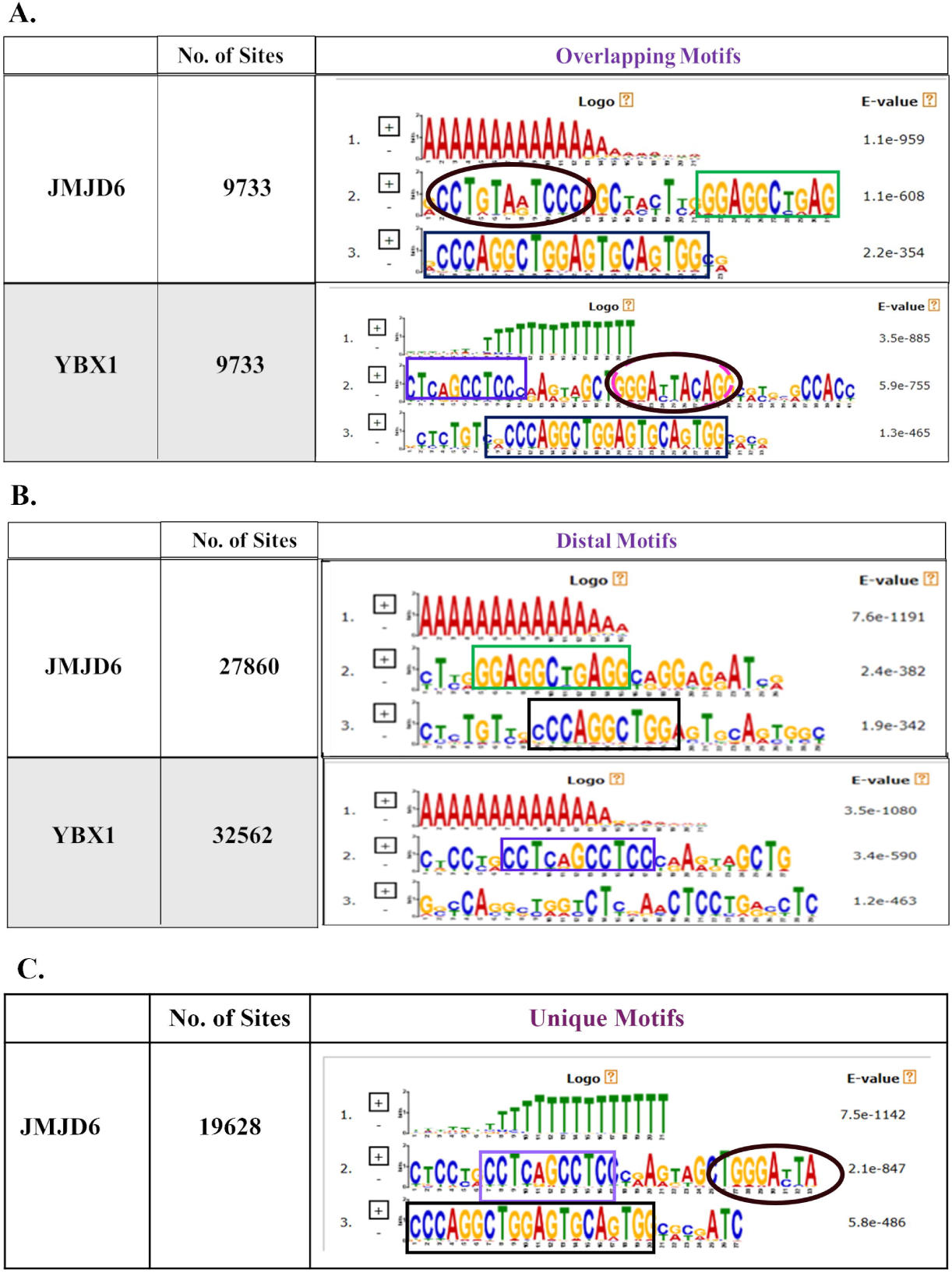
MEME-ChIP analysis to detect JMJD6/YBX1 motifs. A) Motifs detected in common 9733 binding sites B) Motifs found in non-superimposing JMJD6 and YBX1 binding sites C) Motifs of unique JMJD6 binding sites

Independent JBS sites were analysed next and MEME-ChIP and TOMTOM analysis of these sites is shown in Figure 4C. Surprisingly, no unique motif was found but the same 3 motifs identical to the 9733 YBX1 motifs reappeared. In YBX1 ChIP, these sites could remain unreported due to low enrichment that prevented their selection based on statistical cut-offs. This raises the possibility breast cancer cells that express high JMJD6 levels prefer to use YBX1 as their consistent genomic binding partner.

### JMJD6 over expression alters histone marks

JMJD6 is a histone lysyl hydroxylase and known to alter histone marks in the vicinity of its binding (Gao et al., 2015). We estimated enrichment of H3K27Ac (an activation mark), H3K27me3 (a repression mark) and H3K4me1 (an enhancer mark) in JOE cells and compared their genome wide profiles to published parental MCF7 profiles from ENCODE data (Figure 5A). Interestingly, H3K27Ac peaks were abundant near TSS in JOE cells and around 70% of were found within promoter sites (5kb around TSS) while in MCF7 cells only 30% were detected in this region (Figure 5, A and B). Peaks for H3K4me3 showed a major shift from the promoters in MCF7 cells (65%) to distal sites in JOE cells (15%) (Figure 5B).

**Figure 5:**
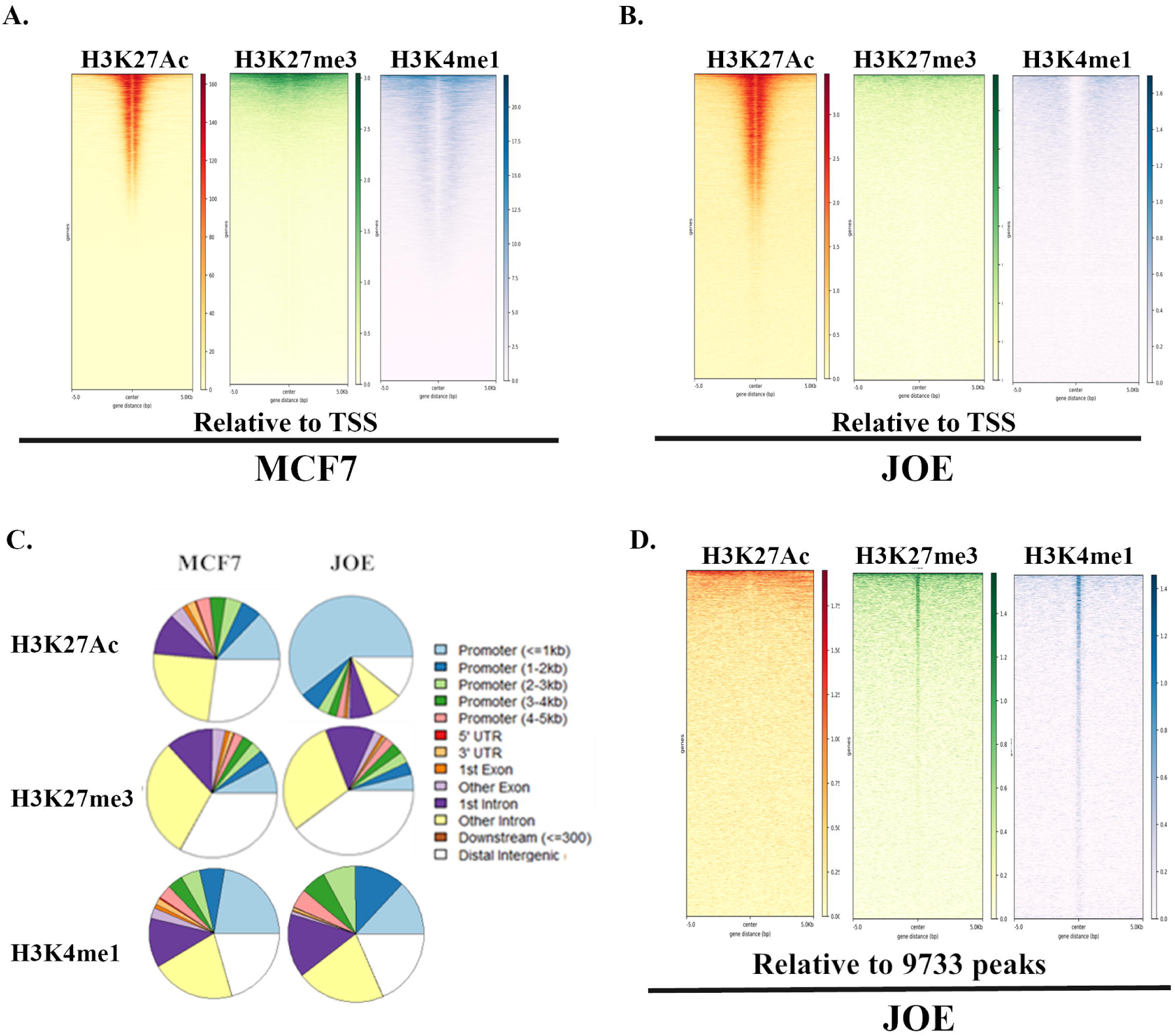
Epigenetic changes driven by high JMJD6. A) Distribution of H3K27Ac, H3K27me3, and H3K27me1 peak distribution surrounding the TSS in A. MCF7 cells; B) JOE cells. C) Pie-charts showing comparison of histone marks based on genomic feature annotation in MCF7 and JOE cells. D. Histone modification landscapes around centered JBS.

These reciprocal changes in histone marks around promoters in JOE cells is indicative of augmented transcription of target genes (Figure 5B). Next, we looked for enhancer mark H3K4me1 and found that its binding was distributed all over the genome and less dramatically altered from MCF7 cells (Figure 5B). Histone modifications in the vicinity of 9733 summit peaks were studied and since the 9733 peaks mostly localized to distal sites, hardly any H3K27Ac peaks were found near the peak summits. Possibly JBS/YBS peaks are away from super enhancer regions. On the other hand, these chromosomal regions had significant enrichment of H3K27me3 and H3K4me1 signals (Figure 5C). These bivalent marks represent poised enhancers during development and lineage determination if present at the same genomic location (Rada-Iglesias et al., 2011). However, in our data, overlap extension analysis of these two modifications showed no overlapping sites, indicating that these marks were not associated with the same site/region and hence did not represent bivalent regions. Further, most of the 9733 peaks consistently associated with H3K4me1 mark suggesting that these peaks were near gene enhancer regions. Together, high expression of JMJD6 appears to promote transcriptionally permissive landscapes allowing elevated gene expression.

### Functional validation of representative BrCa and newly identified sites

We selected multiple common binding sites of JMJD6 and YBX1 for further exploration. We compared the 13 BrCa JBS sites validated in ChIP-reChIp assays (Figure 2) and found that they contained the combination of motifs discovered in this work including the flanking homopolymeric regions. The *AURKB* site was previously validated by us using ChIP-PCR in both MCF7 and MDA MB231 cells as a BrCa JBS (Biswas et al., 2020) and by qRT-PCR for increased RNA expression in this study (Supplementary figure 1). *AURKB* site is also promoter proximal (−353bp near the TSS) to *VAMP2* and this too is an induced gene in our RNA-seq data. This site showed a H3K4me1mark and the promoters display H3K27Ac peaks (Figure 6A). In JMJD6 and YBX1 ChIP experiments both proteins showed enrichment in this region and JMJD6 binding was compromised in YKO cells (Figure 6B). This site was subcloned in pGL4TATA vector (pEBS) and enhancer activity was checked using dual luciferase assays in cells transfected with JMJD6/YBX1 alone or simultaneously. All wells showed enhanced luciferase activity confirming the presence of a regulatory motif in this region (Figure 6C). We tested if deletion of the poly A site and partial YBX1 interacting region (pEBSΔYIR) challenged the enhancer activity. We utilized JOE cells that had a high expression of both JMJD6 and YBX1 with Vec cells as control to perform luciferase assays (Gupta et al., 2025). As shown in Figure 6D, a complete loss of luciferase activity was evident for pEBSΔYIR construct when compared to the intact pEBS construct despite the presence of ample amounts of both proteins in JOE cells. This experiment validated that both YBX1 regions and the homopolymeric tract were essential for JMJD6 mediated enhancement in luciferase activity. Next, we validated two more JBS BrCa sites near *RAD/BRIX* and *IL6* genes. The details of JBS/YBS peaks and histone marks are displayed in Figure 6E. Both sites showed increased luciferase activity when JMJD6 alone or both JMJD6/YBX1 together were transfected in MCF7 cells (Figure 6F). These data indicate that the complete EBS may be essential for optimal regulation of neighbourhood genes. However, if these tracts are required for YBX1 binding at all remains to be seen.

**Figure 6:**
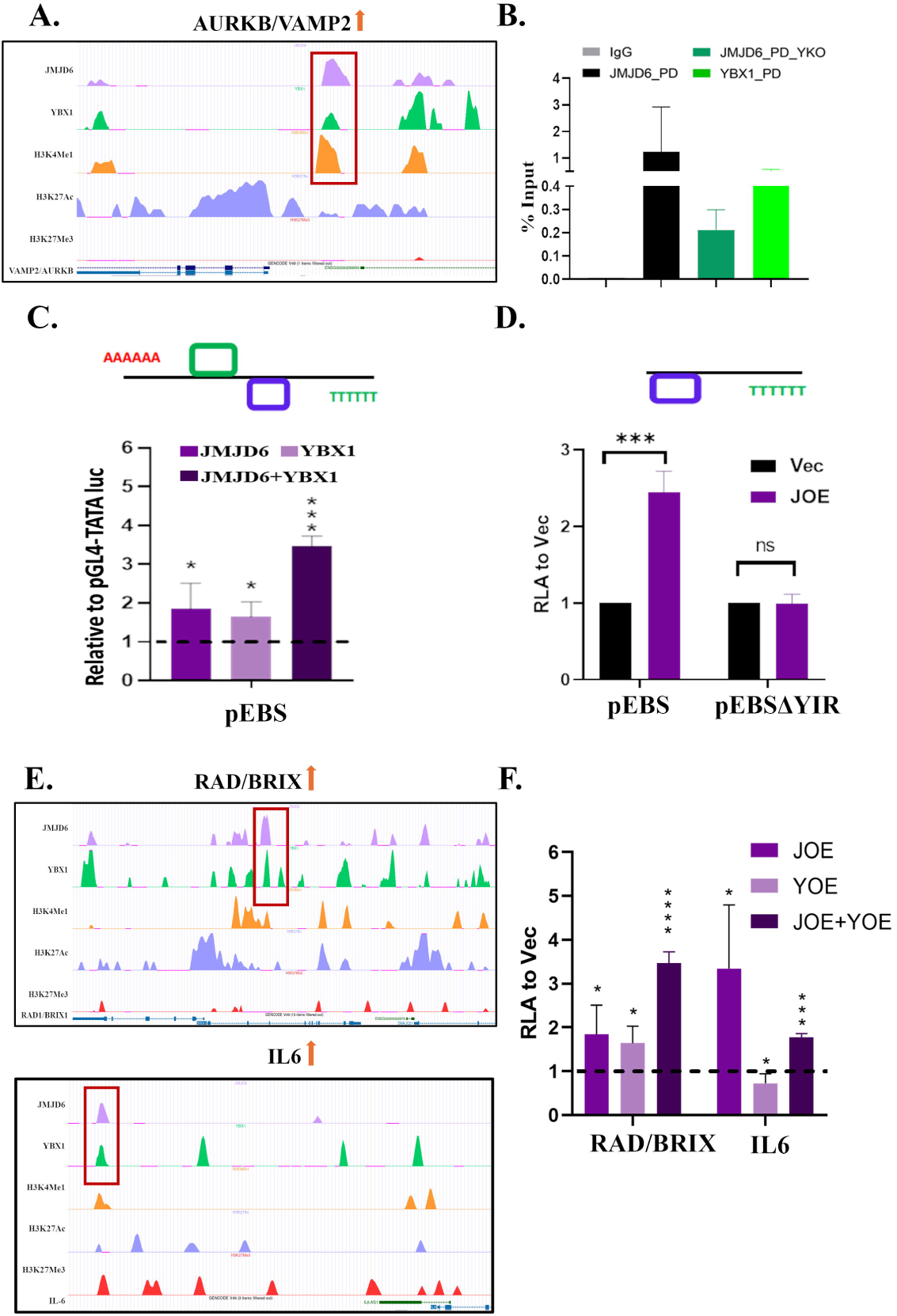
Functional validation of binding sites by enhancer-promoter assays. A) Peak position of JMDJ6 (lavender), YBX1 (green), around VAMP2/AURKB JBS and surrounding histone marks; B) ChIP PCR analysis of JMJD6/YBX1 ChIP DNA and JMJD6 ChIP DNA isolated from YKO cells for VAMP2/AURKB site C) Promoter reporter assay of EBS with overexpression of JMJD6 and YBX1 in MCF7 cells; D) Promoter reporter assay of EBS and YIR deleted EBS in JOE cells; E) Common binding sites of JMJD6 and YBX1 near RAD/BRIX and IL6 genes and surrounding histone marks; F) Promoter reporter assay of RAD/BRIX and IL6 binding sites after over-expression of JMJD6, YBX1 and both proteins.

### Overlap of RNA sequencing and ChIP sequencing data

Post validation of the BrCa JBS sites, we embarked on finding additional genes potentially regulated by JMJD6/YBX1 binding. The peak summits of JBS and YBS sites were mapped to the nearest gene/TSS as described in methods. A total of 17815 different coding/non-coding genes were associated with JBS and 16057 with YBS, of which, 12724 (60%) overlapped (Figure 7A). It was clear that multiple YBX1 sites clustered around JMJD6 sites in regulatory regions of these genes. Of these 12724, 9733 overlapping sites predicted about 6425 annotated genes that could be transcriptional targets of these two proteins. For some genes, more than 2 sites were found. Remainder sites were in the vicinity of un-annotated/predicted genes. However, binding does not always entail differential regulation. To explore the genes differentially regulated by high expression of JMJD6, genes identified in ChIP sequencing data were intersected with DEGs commonly regulated by both proteins from the OE and SI data described above (Supplementary table 3). Overlap yielded 393 (JOE and YOE set) and 198 (Jsi and Ysi) genes (Figure 7B and C). These genes predominantly represented cell proliferation, TGFβ, Interferon and ER targets and E2F regulated pathways. Of interest were Cyclin E2 (CCNE2) and TGFβ2, genes we have previously shown to be responsible for JMJD6 induced proliferation of MCF-7 cells (Lee et al., 2012). Overlapping JBS/YBS peaks were found in the vicinity both genes at (+2000 bp downstream of TSS) for CCNE2 and (−4002bp upstream of TSS) for TGFβ2 (Figure 7D). These were validated by ChIP-PCR. CCNE2 was upregulated in JOE cells and this positive regulation was supported by strong H3K27Ac marks at promoter site. CCNE2 site showed positive enrichment using either JMJD6 or YBX1 ChIP material but enrichment was drastically reduced in YKO cells (Figure 7E). For TGFβ2, the promoter was occupied with H3K27me3, buts no acetylation marks were evident, which probably explains the negative regulation of this gene by JMJD6. Again, both JMJD6 and YBX1 binding was validated in ChIP-PCRs and JMJD6 enrichment was depleted in YKO cells (Figure 7E).

**Figure 7:**
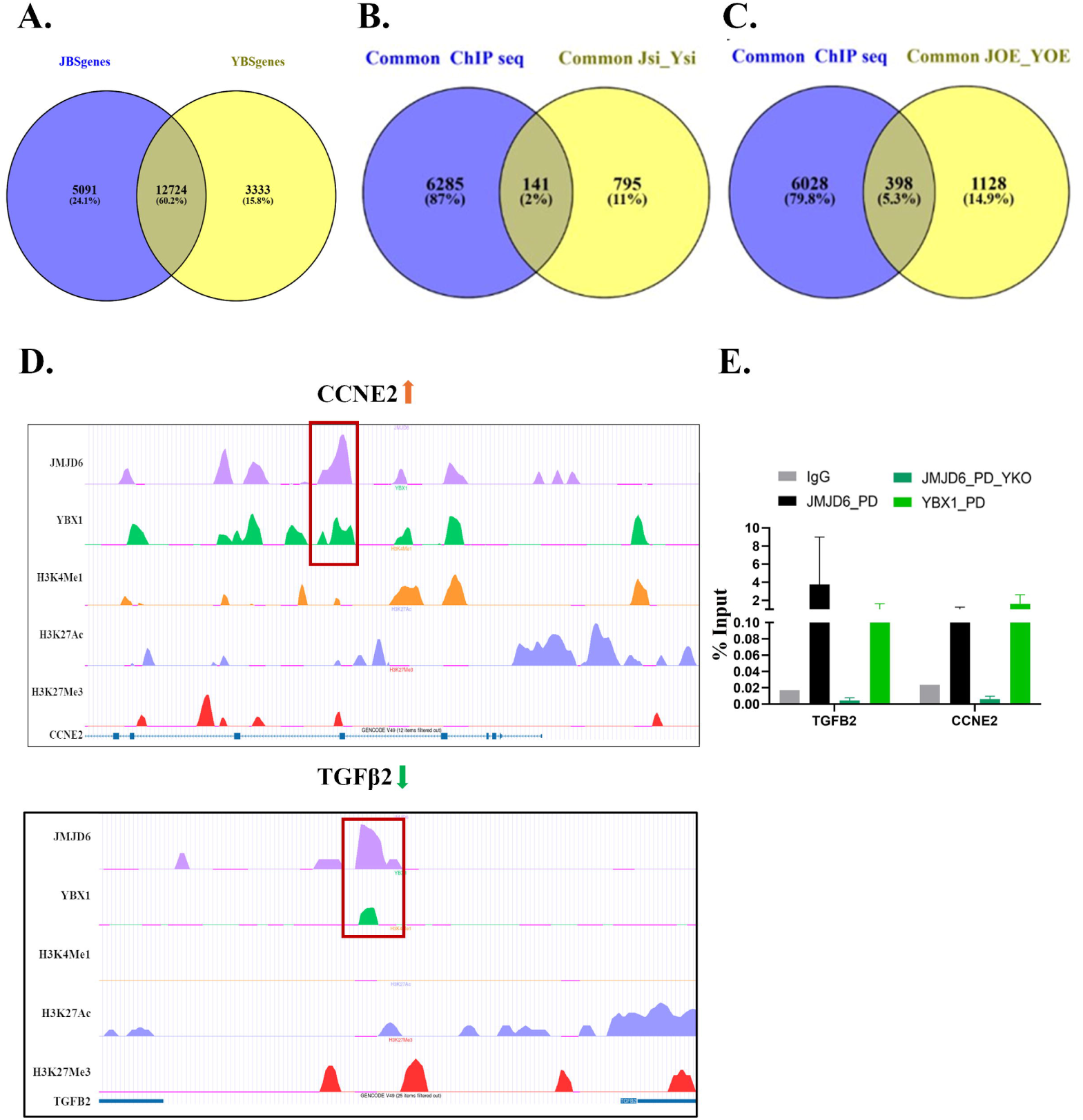
Integration of ChIP sequencing and RNA sequencing data. A) Overlap between genes predicted around JBS and YBS sites; B.-C) Overlap between genes predicted in A with genes commonly regulated by both proteins in RNA-seq data (shown in figure 1); D) Identification of novel overlapping JMJD6 and YBX1 peaks in regulatory regions of known JMJD6 targets *CCNE2* and *TGFB2* genes and landscape of histone marks in respective regions, E) ChIP PCR validation of JMJD6, YBX1 occupancy at identified sites in JOE and YKO cells.

### Inverse relationship of ER with JMJD6 and YBX1

Our lab reported earlier that high JMJD6 led to suppression of ER in MCF7 cells (Das et al., 2022). RT-PCR and RNA-seq data of JOE cells supported that JMJD6 maybe hijacking regulation of those E2/ER targets that sustain breast cancer proliferation and render cells insensitive to endocrine therapy. Other groups have also reported suppression of ER by increased YBX1 expression but at the translational level (Yang et al., 2020). Therefore, we investigated the *ESR1* gene locus for possible binding peaks. ChIP-seq data identified 3 overlapping peaks for JMJD6 and YBX1. Two peaks are upstream of *ESR1* TSS (−5408 bp, - 27377 bp respectively) and one was intergenic (+40862 bp). Only the latter peak appeared in the 9733 overlap list and was characterized by a negative histone mark (H3K27me3) but not associated with the H3K4me1 mark, an organization similar to the *TGF*β*2* locus. Parental MCF7 cells were devoid negative histone marks near the identified JBS/YBS (Figure 8A). Binding of JMJD6 and YBX1 was validated by ChIP-PCR at this location and knocking out YBX1 reduced JMJD6 binding significantly suggesting that low ER may be a result of transcriptional suppression (Figure 8B). JMJD6 does not completely abolish ER expression in our cells, however, YBX1 interacts with ER to negatively regulate its targets and together these two proteins may ensure complete suppression of ER targets (Campbell et al., 2018).

**Figure 8:**
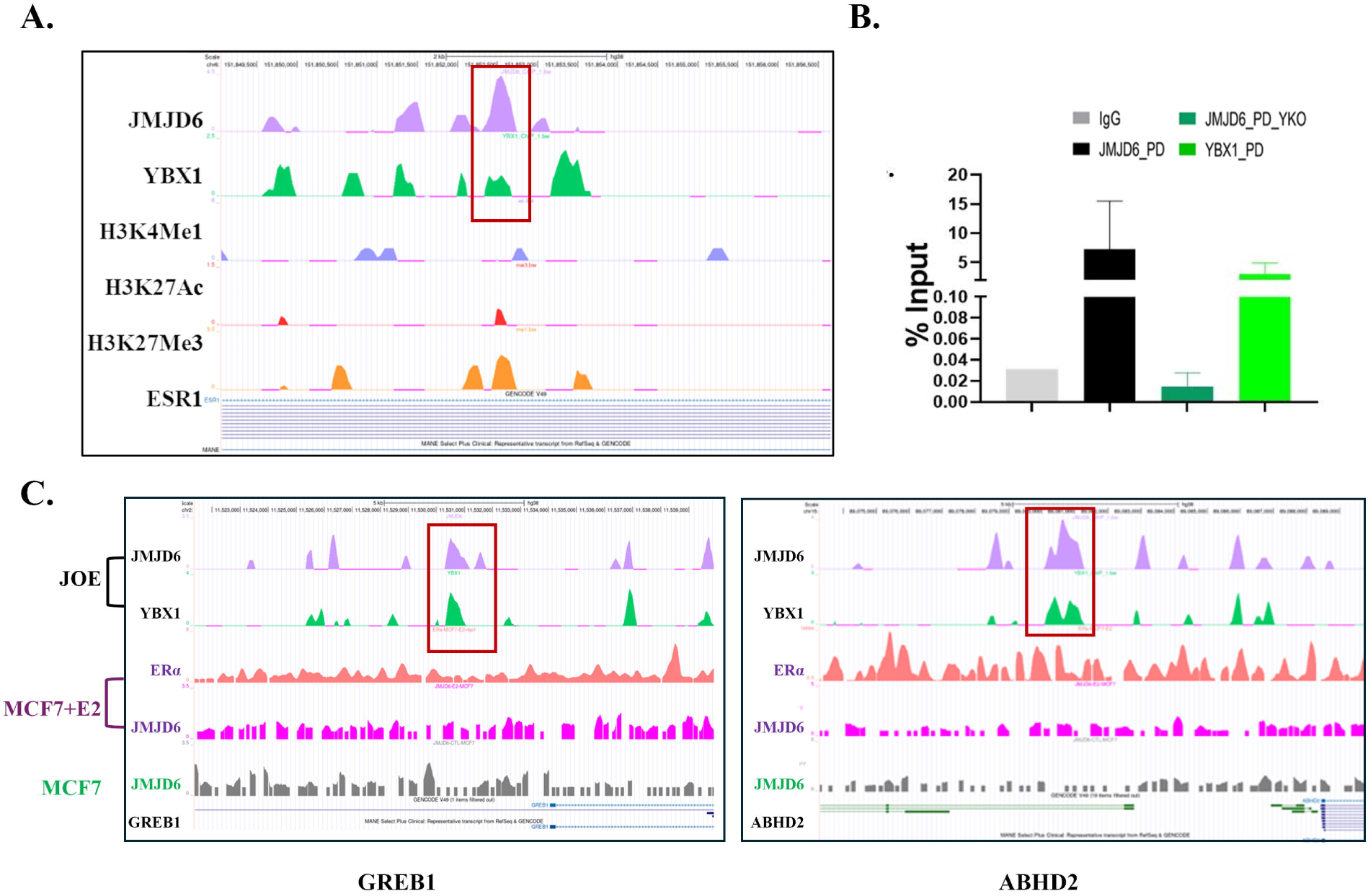
Redistribution of JMJD6 to alternate YBX1 binding sites near *ESR1* genomic locus and representative ER target genes. A) Possible regulatory peaks of overlapping JMJD6 and YBX1 near ESR1 TSS ; B) Validation of common binding site by ChIP PCR in JOE and JOE-YKO cells; C) Binding and recruitment of JMJD6 with the help of YBX1 upstream of TSS of ER regulated genes (ABHD2, GREB1). ER peaks in the presence (deep purple) and absence of estrogen (darker green) and lack of JMJD6 peaks under both conditions are shown.

In contrast, a subset of genes including MYC, E2F and 20/26 ER targets from E2 response pathway, were regulated in a manner similar to an activated E2/ER axis. To determine how this regulation was sustained in absence of E2 and low ER levels in JOE cells, we downloaded publicly available JMJD6 and ER ChIP seq data from MCF7 cells with and without estrogen (E2) treatment. In presence of E2, JMJD6 and ER interaction is essential for RNA pol II pause release to drive the ER transactivation of gene expression program (Gao et al., 2018). Comparison of peak position for ER and JMJD6 in MCF7 cells in presence/absence of estrogen, JBS/YBS in JOE cells and the related histone marks for two representative ER regulated genes ABHD2 and GREB1 is shown in Figure 8B. Clearly, no JMJD6 peaks are found in this region for JMJD6 irrespective of E2 treatment, whereas ER peaks are abundantly enriched following E2 treatment. However, in untreated JOE cells, overlapping JMJD6-YBX1 peaks appear at chromosomal locations other than those of ER, suggesting that regulation may now occur via an alternate site to promote gene expression in JOE cells (Figure 8C). It appears that YBX1 serves as a toggle switch to permit regulation of a subset of ER target genes that are essential for cancer maintenance/progression by carefully recruiting JMJD6 at such sites. Together these data propose mechanism by which JMJD6 sustains E2/ER target gene expression by trading its partnership with ER for YBX1.

### JMJD6, YBX1 expression and correlation with their targets in patient samples

RNA sequencing was performed on 26 patient samples (14 ER+ and 12 ER-). In ER negative samples, both the genes were expressed in higher amounts (Figure 9A). Inter gene comparison showed expression of YBX1 was higher than JMJD6 in patients irrespective of their ER status. In an independent cohort of 22 patients, RT-PCR analyses verified RNA-seq data observations (Figure 9B). These observations could also be extended to publicly available breast cancer TCGA datasets. In Firehose Legacy data (n=1108 samples), similar trends in expression of JMJD6 and YBX1 were noticed (Figure 9C). This re-emphasized that the inverse expression relationship between ER and JMJD6/YBX1 observed in cell line data could be extrapolated to patient samples. To check if differences exist in RNA profiles of representative JMJD6-YBX1 targets defined in this study, gene expression correlation plots for JMJD6, YBX1, ESR1, AR and JMJD6-YBX1 targets AURKinases, CCNE2 and TGFB2 were generated using the KM-plotter tools. Overall, in normal breast, JMJD6 and YBX1 showed negligible correlation to each other, or CCNE2, but maintained inverse relationship to ESR1 and AR. Both AURkinases demonstrated a trend towards positive correlation with these two hormone receptors that typically govern breast cell proliferation, but not JMJD6 nor YBX1. In contrast, JMJD6, YBX1 expression gained positive correlation values between themselves and also to AURkinases, and CCNE2 in breast cancer data. This observation appears to indirectly support our notion that in low ER expressing or ER-cells, JMJD6-YBX1 axis may take over the regulation of cell proliferation and cell division.

**Figure 9:**
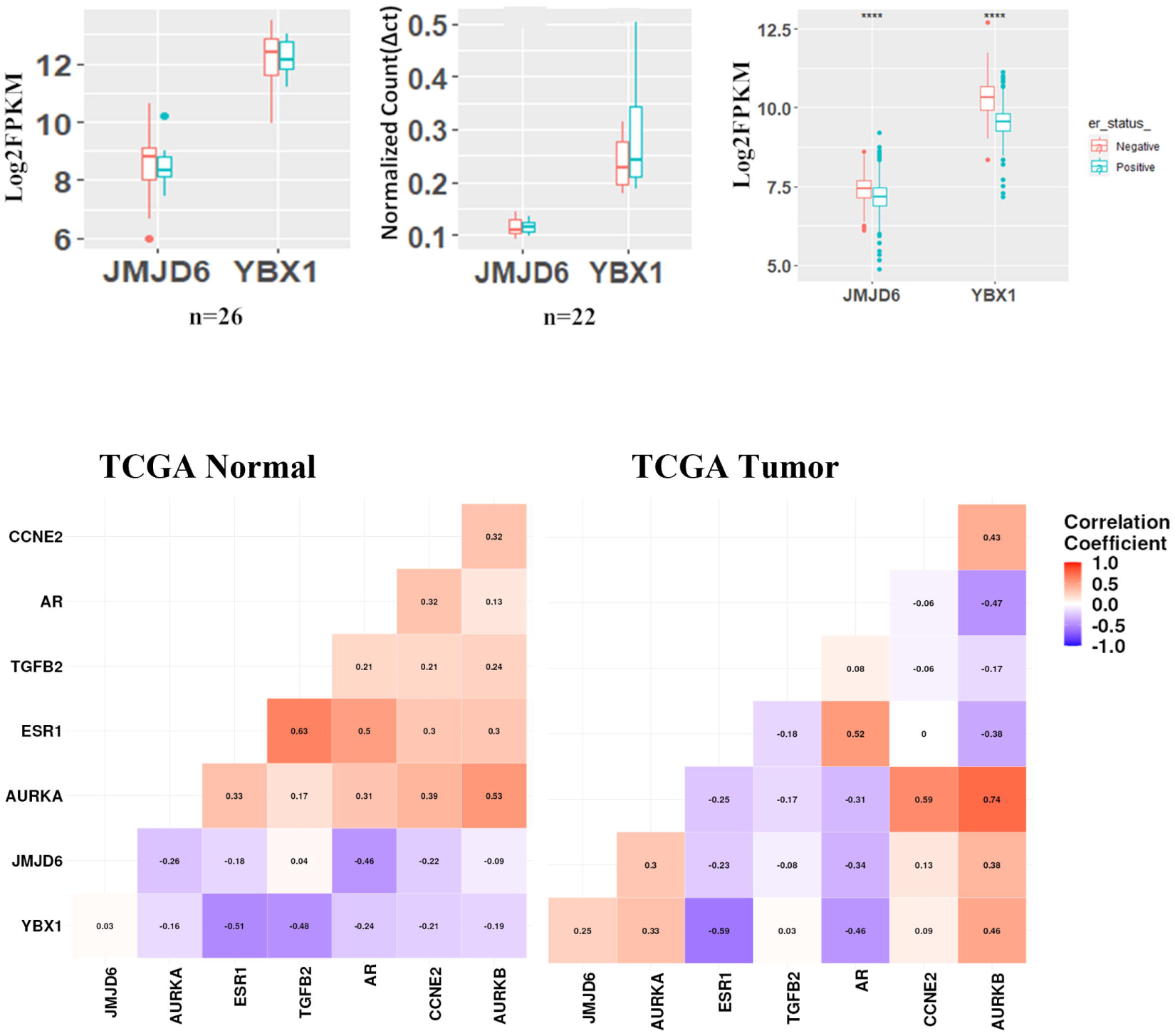
Expression of JMJD6 and YBX1 in patient samples. A) Expression of JMJD6 and YBX1 from RNA sequencing of 26 Breast Cancer patients; B) qRT-PCR of JMJD6 and YBX1 in a different cohort of patients (n=22); C) RNA expression of JMJD6 and YBX1 in TCGA Firehose Legacy data; D) Gene correlation analysis in TCGA normal and TCGA breast cancer RNA-seq data

## Discussion

Prior studies have independently indicated that both YBX1 and JMJD6 physically interact with ER following estrogen stimulation and improve its transcriptional activity under physiological conditions (Campbell et al., 2018; Poulard et al., 2014). Worse survival, cancer progression and resistance to endocrine therapy was indicated in patients who showed enhanced expression of either protein (Bargou et al., 1997; Lee et al., 2012). Studies in breast cancer cells have suggested that the probable mechanism could be JMJD6/YBX1 mediated depletion of ER expression (Campbell et al., 2018; Das et al., 2022). Here we confirm that YBX1 recruitment of JMJD6 and change in histone modification landscape at the ESR1 locus mediates its negative transcriptional regulation. Our earlier studies showed that high levels of JMJD6 compensated for lack of ER by positively regulating ER-targets that were critical for cancer maintenance possibly driving endocrine therapy resistance. Recently, our work demonstrated that JMJD6 physically interacts with YBX1, they enhance each other’s expression and YBX1 was critical for JMJD6 recruitment to regulate the proximal promoter of the oncogenic lincRNA HOTAIR. Here we extend these ideas to explore if YBX1 partners with JMJD6 binding in a genome wide manner and if this results in co-regulation of cancer promoting genes under conditions of low ER expression.

Transcriptomic analyses revealed extensive overlap in JMJD6- and YBX1-regulated gene expression, both in gain- and loss-of-function perturbations. Shared DEGs predominantly mapped to estrogen-response pathways, cell-cycle progression (E2F, G2-M), interferon programs and immune-regulatory modules, all events critical to cancer progression. In addition, our results propose a complex regulatory interplay involving JMJD6 and YBX1, siRNA knockdown induced a subset of genes with very large fold-changes, yet only a narrow range of genes were significantly repressed. Conversely, JMJD6/YBX1 overexpression led to a broad suppression of many genes (some with substantial downregulation), while the set of upregulated targets was limited with modest increases. This asymmetry suggests that JMJD6 and YBX1 may not act as straightforward on/off switches for the same genes but rather have distinct influences depending on context. One plausible explanation is that these proteins carry out independent post-transcriptional functions such as mRNA stability control. For example, YBX1 overexpression can prolong mRNA half-lives by recruiting RNA stability factors such as ELAVL1, whereas YBX1 depletion accelerates transcript decay (Miwa et al., 2006; Sánchez et al., 2023). X-ray crystallographic studies have projected an RNA binding function for JMJD6 that is currently being explored in various studies (Hong et al., 2010). Possibly, JMJD6–YBX1 complexes target specific transcripts for degradation or protection, removing these proteins and conversely excessive proteins cause imbalances in transcript amounts supporting pathological outcomes. These ideas warrant further detailed experimental validation. Of note is the known involvement of both proteins in alternative splicing, expanding their role further in post-transcriptional regulation and RNA homeostasis.

Our focus in this manuscript was to uncover transcriptional programs under JMJD6 and YBX1 control. JMJD6 is not known to bind DNA directly and historically has been considered a secondary chromatin interactor that is guided by transcription factors. Our analyses now identify YBX1 as the predominant DNA-binding partner that specifies the genomic distribution of JMJD6. Through proximity ligation, ChIP-PCR, ChIP-reChIP assays, KO-based ablation experiments, and genome-wide profiling, we demonstrate that YBX1 is required for JMJD6 enrichment at atleast 13 validated breast-cancer JMJD6 binding sites. Importantly, transient knockout of YBX1 resulted in a loss of JMJD6 occupancy at all 13 BrCa JBS sites, strongly arguing that YBX1 is indispensable for JMJD6 chromatin loading. Genome-wide ChIP-seq profiling revealed that ∼20% of JMJD6 and YBX1 peaks were physically superimposed, and nearly all remaining JMJD6 peaks contained a neighbouring YBX1 site within ±50 kb, consistent with looping-based enhancer–promoter communication. These patterns suggest that JMJD6 is rarely, if ever, positioned on chromatin independently; instead, YBX1 acts as a scaffold, recruiting JMJD6 across the genome to enhancers and promoter-proximal sites. Motif analyses uncovered an unexpected but highly consistent architecture: (i) flanking poly-A/poly-T tracts; (ii) a composite motif containing the canonical YBX1 recognition sequence; and (iii) a second complementary motif enriched for binding signatures of zinc-finger proteins such as ZNF460 and ZNF213. The convergence of these motifs at 73–80% of JMJD6–YBX1 co-bound sites suggests the presence of a larger multi-protein transcriptional complex. This “Extended Binding Site” (EBS) motif was also present in previously validated JMJD6 sites discovered in HeLa cells and validated by ChIP-PCR in breast cancer cells, indicating that JMJD6–YBX1 genomic interactions follow a conserved sequence logic (Biswas et al., 2020).

Interestingly, ER itself is a Zn-finger transcription factor that interacts with both JMJD6 and YBX1. Discovery of additional Zn-finger binding motifs in JBS/YBS sites may reveal a broader propensity of these factors to prefer this family of proteins as partners for gene regulation. This implicates a kind of conceptual gene regulatory ‘degeneracy’ that governs transcription of key genes when the cells encounter changing physiological conditions such a development and differentiation. However, the same is exploited in cancer, by merely trading one Zn-finger protein for another to sustain dysregulated gene expression, when the original factor is compromised (herein ER).

The importance of the ‘extension’ is revealed by our luciferase reporter assay studies. Functionally, deletion of the poly-A tract and partial YBX1-interacting region from the AURKB/VAMP2 enhancer abolished luciferase reporter activity, even in JMJD6-overexpressing cells. Additional enhancer–promoter reporter validations at RAD/BRIX and IL6 loci further confirmed that intact EBS architecture is indispensable for transcriptional activation. Taken together, these findings define a sequence-encoded grammar for JMJD6–YBX1 enhancer regulation.

Simple homopolymeric repeats are not canonical transcription factor binding sites but are found enriched at boundaries of chromatin regulatory regions. Prior studies show that they could play a multifaceted role: (i) Recruiting factors to DNA regions like TBP at TATA-less promoters. Homopolymeric T stretches in promoter regions are known to substitute for TATA boxes in a significant fraction of gene and TBP binds with high affinity to these stretches (Rada-Iglesias et al., 2011). TBP also interacts with YBX1 (Eliseeva et al., 2011). Though, very few JBS/YBS sites were located in promoter regions, the possibility of long-range interactions of JMJD6/YBX1 to promoter regions cannot be ruled out. More interesting though, is the recent finding that homopolymeric DNA can recruit specific RNA-binding coactivators such as PABPC1 that ordinarily interact with polyA stretches in RNA. This factor was shown to bind polyA DNA and co-occupy ETS regulatory regions in prostate cancer (Wigington et al., 2014). PABPC1 appears as a potential interactor in mass spectrometry data generated to find JMJD6 partners and PABPC1 is known to bind YBX1 (Zheng et al., 2025). This represents a non-traditional mode of gene regulation where an RNA-binding protein contributes to DNA-based transcriptional complexes. (ii) Imposing pausing checkpoints in transcription elongation as seen in *MYB* transcripts. This mRNA harbors a poly-T sequence downstream of TSS where RNA pol II stalls and transcription elongation is halted. The elongation can continue only after an E2/ER dependent release. ER binding subsequently recruits the positive transcription elongation factor b (P-TEFb) – a kinase complex of CDK9 and cyclin T1 (Mitra, 2018). This factor is known to interact with JMJD6 for its anti-pause release function in transcription initiation (Liu et al., 2013; Zhao et al., 2015). This raises the interestingly possibility that JMJD6/YBX1 might also compensate for ER regulatory function in transcriptional elongation since almost 40% of JBS/YBS sites were located in the gene body. (iii) Creating an accessible chromatin environment: Poly(A/T) tracts are well-known to be intrinsically stiff DNA sequences that resist bending. As a consequence, they disfavor nucleosome formation and tend to exclude nucleosomes and create nucleosome-depleted regions (NDRs) (Field et al., 2008). Studies in yeast and human have shown that long poly(dA:dT) tracts cause large local nucleosome depletion not only over the sequence itself but also extending into flanking regions. This yields an open chromatin configuration and enhances accessibility for transcription factors at nearby binding sites. The presence of a poly(dA:dT) “minisatellite” could serve as a boundary element that partitions chromatin into an open domain, thereby facilitating the loading of transcriptional machinery or co-factors that JMJD6/YBX1 recruit.

Certainly, there is substantial change in accessible chromatin environment since JMJD6 overexpression significantly reconfigures histone landscapes in a genome-wide manner. Promoter-proximal H3K27ac marks were markedly increased in JOE cells, whereas the repressive H3K27me3 mark was globally depleted from promoters, consistent with enhanced transcriptional activation. Conversely, enhancer-associated H3K4me1 peaks showed broad, genome-wide distribution with only modest changes, suggesting that JMJD6’s major epigenetic impact may occur at promoters through acetylation-associated activation, and potentially via histone lysyl-hydroxylation-mediated chromatin remodeling. However, though JMJD6 is well characterized histone-lysyl hydroxylase, we were unable to establish specific pull-downs for this modification using commercially available antibodies in our lab.

Interestingly, at the 9733 co-bound distal enhancer sites, H3K27ac was largely absent, but H3K27me3 and H3K4me1 signals were enriched, though at non-overlapping locations. The former indicating poised or developmentally regulated enhancer states and latter showing co-occurrence of 9733 sites at enhancer loci. These observations highlight that JMJD6–YBX1 complexes act to reshape histone landscapes in a manner that amplifies transcriptional output from growth-related gene networks. Overall, these features align well with the dynamic regulatory functions ascribed to JMJD6 and YBX1 in cancer cells, where precise control of gene expression – at the levels of chromatin openness, transcription initiation, elongation, and mRNA stability – can contribute to cellular transformation and tumor progression. This functional versatility could be attributed to their capability to bind both DNA and RNA.

Next, integration of ChIP-seq and RNA-seq datasets was conducted uncovering 393 upregulated genes in JMJD6- and YBX1-overexpressing models and 198 downregulated genes upon siRNA-mediated depletion, representing high-confidence functional targets. This is an under-representation of regulated genes since programs like ChIPseeker only assign the TSS of ONE nearest gene to any identified site. Nevertheless, this sub-set included CCNE2 and TGFβ2, genes previously implicated in JMJD6-driven proliferation (Lee et al., 2012). Both loci harboured JMJD6–YBX1 peaks, showed histone signatures consistent with activation and repression respectively, and displayed reduced JMJD6 occupancy upon YBX1 knockout. This strongly supports a model wherein YBX1 recruits JMJD6 to regulate genes. Perhaps the most compelling insight from this study is that the JMJD6–YBX1 axis can sustain estrogen-response gene expression independently of ER. Importantly, 20 of 26 canonical ER target genes were regulated concordantly by JMJD6 and YBX1, indicating that the JMJD6–YBX1 axis sustains core ER-dependent transcriptional circuits. ER loss in JMJD6-overexpressing cells was accompanied by the emergence of JMJD6–YBX1 enhancer sites at alternate genomic positions relative to canonical ER–JMJD6 sites described in estrogen-stimulated cells. This was particularly evident at multiple locations such as ABHD2 and GREB1, where JMJD6–YBX1 binding replaced ER-dependent regulatory engagement. Interestingly, peaks for these two factors were also present in the vicinity of ESR1 locus without corresponding H3K4me1 peaks indicating that suppression of ESR1 locus perhaps occurs at the level of transcription. These findings offer a mechanistic explanation for earlier reports from our group and others showing that high JMJD6 or YBX1 suppresses ER expression to promote endocrine therapy resistance.

Consequently, we propose a model wherein JMJD6 and YBX1 switch their transcriptional partner from ER to other ZNF family members using predominantly YBX1 mediated JMJD6 recruitment to YBX1 DNA binding motifs instead of EREs, when JMJD6 levels are aberrantly elevated (Figure 9). This rewired enhancer logic enables breast cancer cells to maintain proliferation-driving ER target gene expression in an ER-independent manner, contributing to endocrine unresponsiveness and disease progression. Under physiological conditions however, JMJD6 and YBX1 assist the master regulator E2/ER mediated transcription to maintain breast physiology.

Finally, to assess the real-world tumor sample data for JMJD6 and YBX1, patient-derived RNA-seq, RT-PCR analysis from independent cohorts were employed. They consistently showed elevated JMJD6 and YBX1 expression in ER-negative tumors, and an inverse correlation between ER and JMJD6/YBX1 in ER+ tumors. TCGA datasets recapitulated this trend across >1,000 samples. These data suggest that the JMJD6–YBX1 transcriptional axis may be preferentially activated in more aggressive, ER-low or ER-negative breast cancers. Given that both proteins individually associate with poor prognosis and therapeutic resistance, their cooperative function revealed here suggests that JMJD6–YBX1 may serve as a powerful biomarker pair and a potential therapeutic target. Inhibiting their interaction, disrupting EBS architecture, or blocking JMJD6 enzymatic activity could provide new avenues to overcome endocrine therapy resistance.

## Conclusion and Future Directions

Our study uncovers a fundamental mechanism by which breast cancer cells escape ER dependence: the hijacking of ER enhancer networks through a JMJD6–YBX1 transcriptional module. By defining the sequence grammar of JMJD6–YBX1 binding, demonstrating their cooperative recruitment at enhancers, and establishing their ability to sustain key ER-regulated pathways independently of ER, we reveal a new paradigm of transcriptional rewiring in breast cancer progression. Targeting JMJD6–YBX1 interactions, modifying enhancer accessibility, or inhibiting JMJD6 enzymatic activity represent promising therapeutic strategies to counter endocrine resistance. Future studies integrating chromatin topology, single-cell analysis, and *in vivo* models will be crucial to translate these findings toward clinical intervention.

## Supplementary figures

**Supplementary figure 1:** A) RT-PCR analysis of selected genes from RNA-seq data. Constructs transfected and gene names are indicated below the graph. B) RT-PCR validation of select genes from the siRNA treatment RNA-seq dataset. C) Immunoblots showing induction or suppression of genes following recombinant expression or siRNA treatment respectively.

**Supplementary figure 2:** Presence of potential ZNF motifs in EBS.

**Supplementary figure 3:** A) Co-expression of JMJD6 and YBX1 in TNBC/ER-samples in TCGA data. B) Heatmap showing DEGs and the 5 most enriched GO pathways in JMJD6si and YBX1si treated MDA MB 231 cells. C) Enrichment of JMJD6 on BrCa JBS sites and YBX1 on JBS sites in MDA MB 231 cells

## Supplementary tables

**Supplementary table 1:** Primers used in this study

**Supplementary table 2:** Genes regulated in OE and siRNA treated MCF7 cells

## Declarations

Ethics Approval and Consent to participate-Human samples were used in this study were approved by the Institutional review board (NIBMG/20/2018-19)

## Availability of data and materials-

Transcriptomic and ChIP-seq data generated in this paper is deposited in GEO omnibus. (GSE211031 and GSE296699 respectively)

## Authors contributions

A Gupta: Data generation, visualization, analysis. S Bhardwaj: Data generation, visualization and curation. KV Desai: Conceptualization, Supervision, Project coordination, data generation, analysis. All authors contributed to writing of first draft, review and editing of final draft of manuscript.

## Acknowledgement

NIBMG genomics core facility is acknowledged for extending use of their instrumental facility.

