## Supplementary Table 1 for "Genome wide analysis of YBX1-mediated redistribution of JMJD6 in ER+ breast cancer cells"

Table 1: Primers used in this study

| **Guide RNA sequences** | |
| --- | --- |
| **Name** | **Sequence (5'-3')** |
| JMJD6_sg_RNA | ACCGTCTTGTGCCATACCAC |
| YBX1_sg_RNA | GTGTACAAATACATCTTCCT |
| **qRT PCR primers** | |
| JMJD6-RT-F | GGT TGA CCT TCA GGA GTC CAC |
| JMJD6-RT-R | TGC GCT CTT TGC TGA CAC AGT C |
| YBX1-RT-F | GCAGGAGAACAAGGTAGACCAG |
| YBX1-RT-R | CTTCATTGCCGTCCTCTCTAGG |
| Beta-actin-F | GAG CAC AGA GCC TCG CCT TT |
| Beta-actin-R | TCA TCA TCC ATG GTG AGC TGG |
| HOTAIR-RT-F | GGTAGAAAAAGCAACCACGAAGC |
| HOTAIR-RT-R | ACATAAACCTCTGTCTGTGAGTGCC |
| SIRT4-RT-F | GTGGATGCTTTGCACACCAAGG |
| SIRT4-RT-R | GGTTCAGGACTTGGAAACGCTC |
| ADAM17 RT-F | AACAGCGACTGCACGTTGAAGG |
| ADAM17 RT-R | CTGTGCAGTAGGACACGCCTTT |
| AURKA RT-F | GCAACCAGTGTACCTCATCCTG |
| AURKA RT-R | AAGTCTTCCAAAGCCCACTGCC |
| AURKB-RT-F | GGAGTGCTTTGCTATGAGCTGC |
| AURKB-RT-R | GAGCAGTTTGGAGATGAGGTCC |
| RAD1 RT-F | GGATATGACGAGTGAAGTCCTAC |
| RAD1 RT-R | AGTCAAGGTGGGAACTTCCTGC |
| BRIX RT-F | CCAACCATTTGTGGACCACGTG |
| BRIX RT-R | TGGTCCTCCAAAACTTCCCTGG |
| PARP RT-F | CCAAGCCAGTTCAGGACCTCAT |
| PARP RT-R | GGATCTGCCTTTTGCTCAGCTTC |
| NDRG RT-F | ATCACCCAGCACTTTGCCGTCT |
| NDRG RT-R | GACTCCAGGAAGCATTTCAGCC |
| IGF2BP3 RT-F | TCGTGACCAGACACCTGATGAG |
| IGF2BP3 RT-R | GGTGCTGCTTTACCTGAGTCAG |
| IDE RT-F | TACCTCCGCTTGCTGATGACTG |
| IDE RT-R | ACAGGAGCTGAGGTATGAAGGC |
| CISH RT-F | GCATAGCCAAGACCTTCTCCTAC |
| CISH RT-R | ACGTGCCTTCTGGCATCTTCTG |
| **ChIP primers** | |
| HOTAIR-F-216 | CGAGCTCGAAAAGAGAGGGGTGGGAAGG |
| HOTAIR-R-50 | CCGCTCGAGAGTCCTCACTGTGGAAGCTTT |
| MET-1-F | TTGACCTTCACACACCCAGAT |
| MET-1-R | TTCTGAGTTTGAGTGCCATGA |
| SOX2A-F | GAGAGAAA AAGGAGAACCTTCG |
| SOX2A-R | ACGGTGCATTG TTTTGTTCC |
| IL6-F | CAGCTCACTGCAACTTCTGC |
| IL6-R | GAGATCAGCCTGGCAAACAT |
| IGF2BP3-F | GCCTAGGCGACAAGAGTGTT |
| IGF2BP3-R | TGACAAAACTGCTGACAAGCG |
| SIRT4-F | GGCTCAGTGCAGCCTAAATC |
| SIRT4-R | AAAATTAGCTGGGCATGGTG |
| SLCOC3A-F | GGCCTGAGATCCTGCTCAAA |
| SLCOC3A-R | GTGGGGTGGGAGTCTCAATC |
| VAV1-F | TGGCAGGACCAATCACCTTC |
| VAV1-R | CAGCTGGGAATGTCGTGAGT |
| AURKA-F | AGGCTGTGAGATGGCTTGAG |
| AURKA-R | CGATTCTCATGCCTCAGCTT |
| AURKB-F | CCCGCCTCACGATTAAGGTT |
| AURKB-R | CCCGCCTCACGATTAAGGTT |
| ADAM17-F | GGCCAGATCCTGTCTCAAAA |
| ADAM17-R | CAGCCTCCCAAGTAGCTGAG |
| RAD&BRIX-F | TAGCCGTACTAGTTTCAGCCAG |
| RAD&BRIX-R | GTCTTATTGGAGAACTGACCCCT |
| DNAJC21-F | TCAGACAGGTAGAAGGTTTTCG |
| DNAJC21-R | GTGGTGGCACACACCTGTAG |
| IDE-F | GGTCAGGAGTTGGAGACCAG |
| IDE-R | CAACCCCTGTCTCTCGTGTT |
| PARP-F | GAAGGGCTGGCTGCATTA |
| PARP-R | TATGCAGGTGAAGGAAATCAAA |
| NDRG1-F | CCACGCTGAAGACCTCAGTT |
| NDRG1-R | GGCTCAGCTCACAGGTCTTT |
| CISH-F | GTCACAGCCAAGAGCTCAGT |
| CISH-R | CAGGTGAGACAGGCCAGATG |
| NUP1-F | GGGGTACCGCCTCCTCCCACCACTAAC |
| NUP1-R | CGC CTAGGTGAGAGCATGGGTAAAAGG |
| **pEBS Deletion construct Primers** | |
| AURKB_del_EBS_F | CCCGCCTCACGATTAAGGTTC |
| AURKB_del_EBS_R | TGCTGACTGGCCAAGACCTAG |
