## Supplementary figures and images for "Genome wide analysis of YBX1-mediated redistribution of JMJD6 in ER+ breast cancer cells"

### Supplementary figure 1

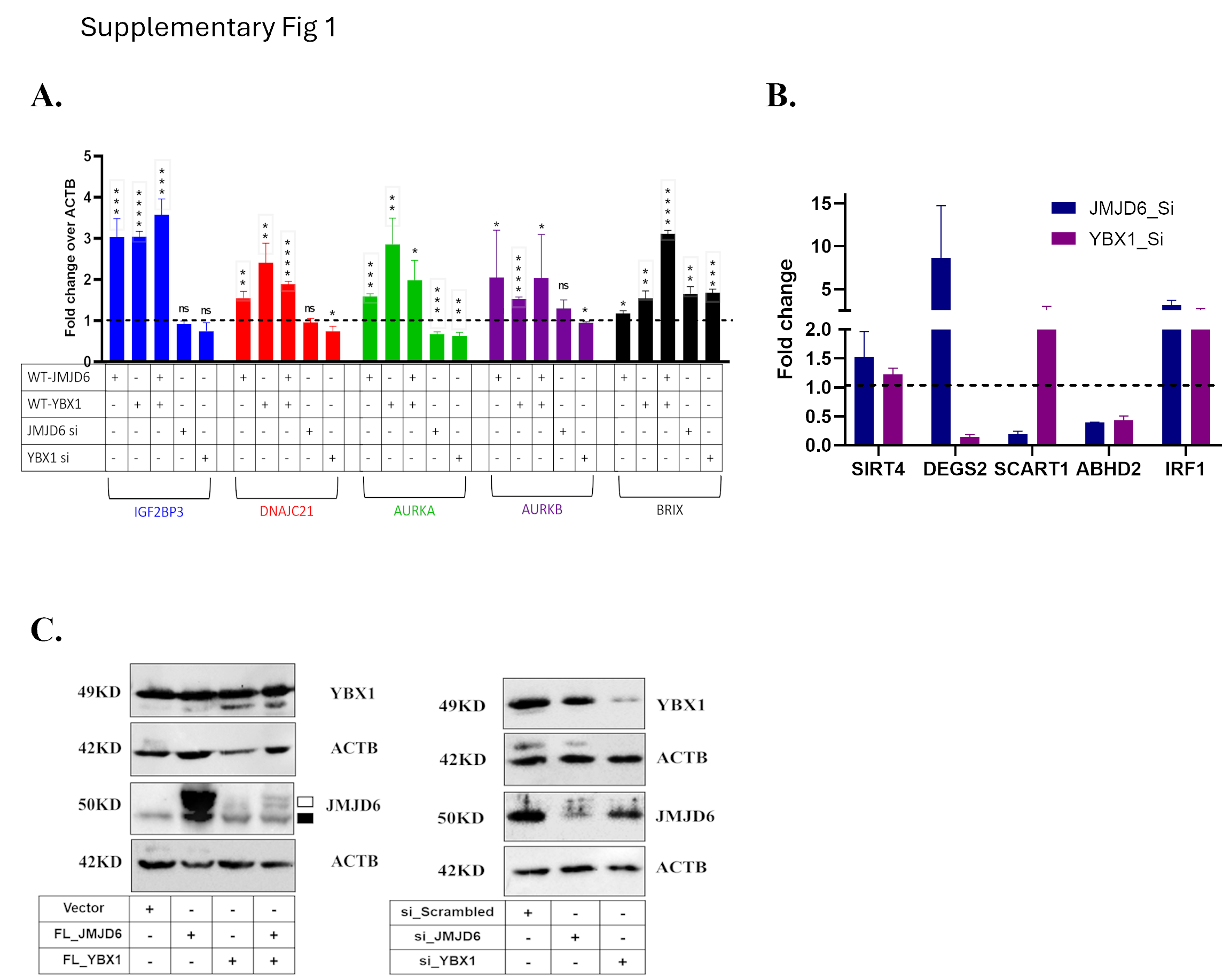

### Supplementary figure 2

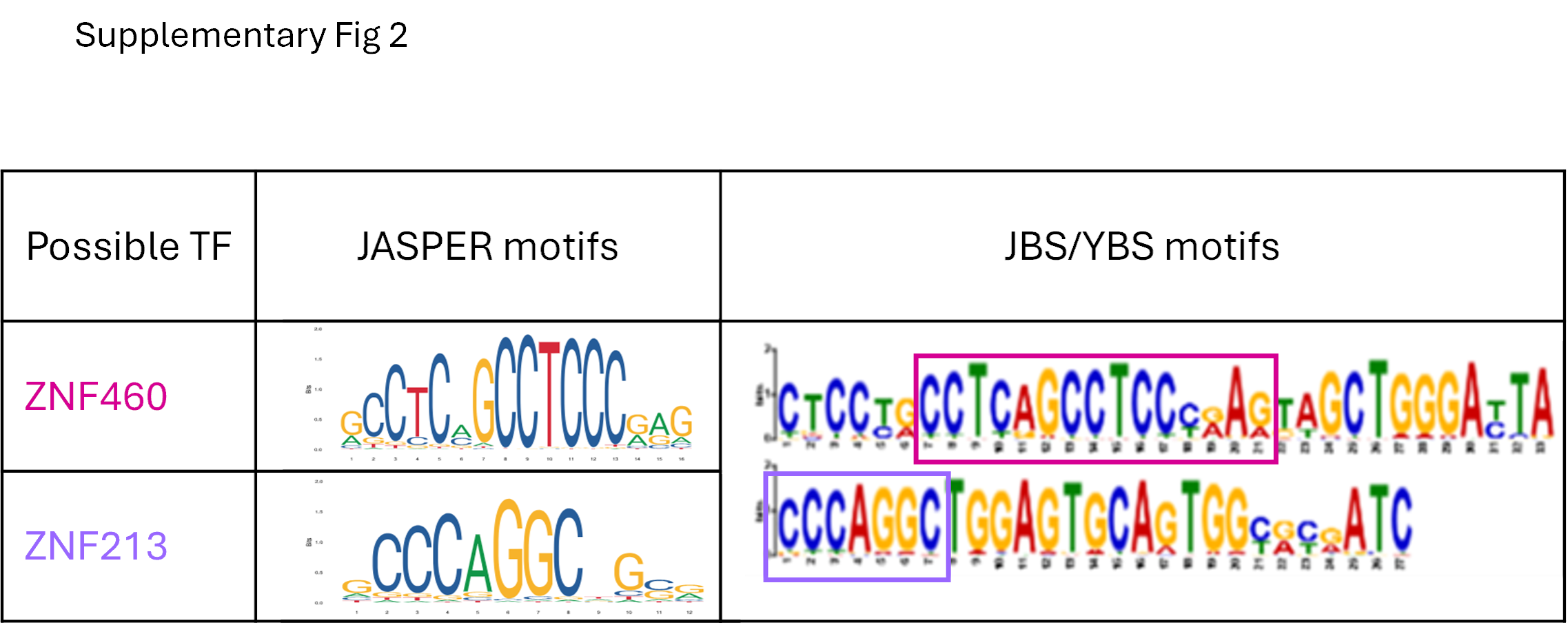

### Supplementary figure 3

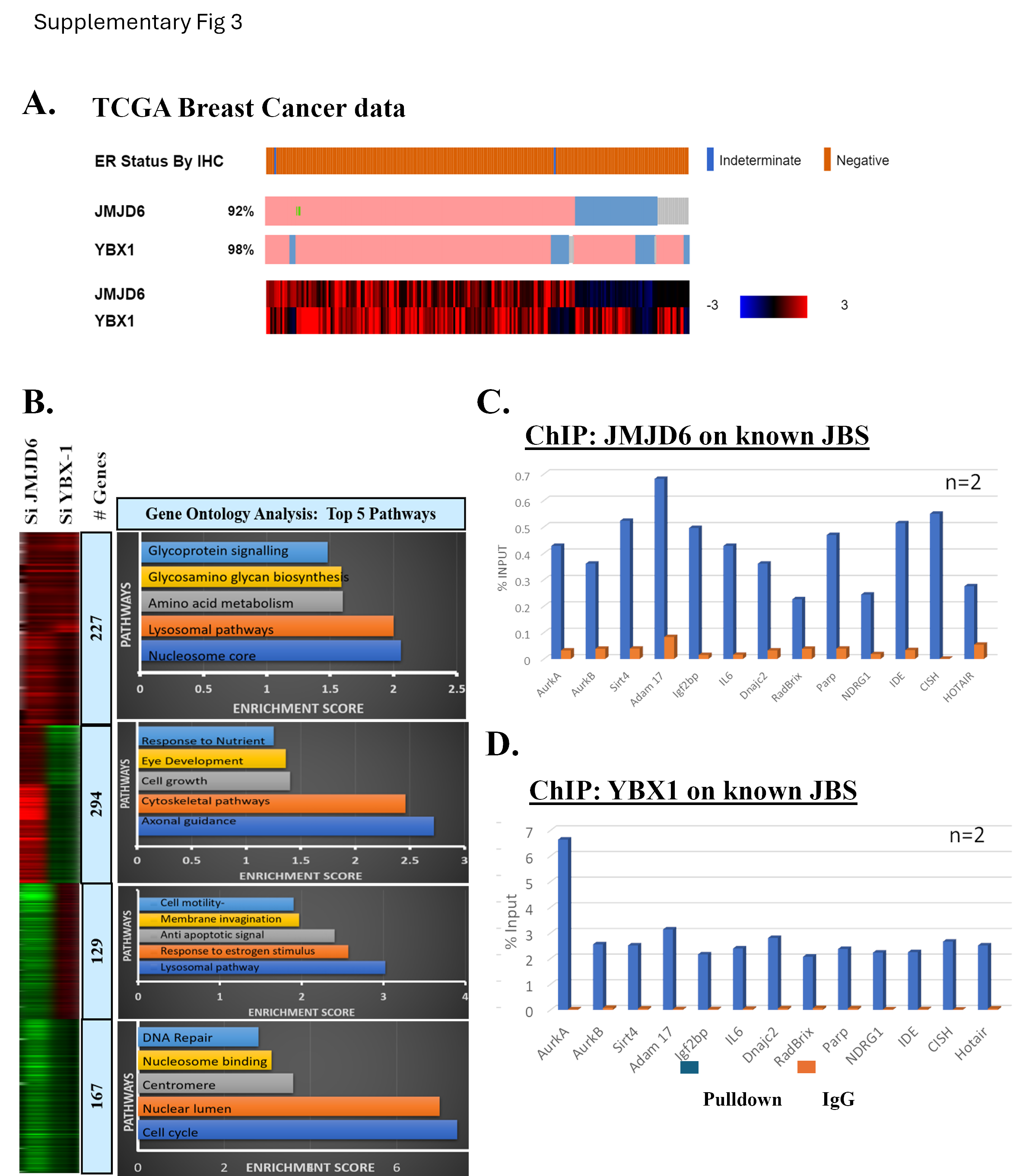
